# Role of TbTim17 in mitochondrial stress tolerance in *Trypanosoma brucei*

**DOI:** 10.64898/2026.06.12.731992

**Authors:** Minu Chaudhuri, Railyn Webster, Hira Karim

## Abstract

Mitochondrial protein translocases Tim17 and Tim23 play important roles in stress-response pathways via activating the transcription factors, ATFS-1, to maintain organellar homeostasis. *Trypanosoma brucei*, a divergent eukaryote and the infectious agent for African trypanosomiasis, lacks ATFs but possesses TbTim17, an essential component of the TIM complex in mitochondria. However, it has not been investigated whether TbTim17 plays a role in the mitochondrial stress response. Here, we show that depletion of TbTim17 increased *T. brucei* tolerance to paraquat, increasing the EC50 by 3- to 4-fold compared with the wild type. Subsequent analysis revealed that increased levels of mitochondrial reactive oxygen species resulting from TbTim17 knockdown upregulate mitochondrial superoxide dismutase, thereby preadapting cells to resist paraquat-induced oxidative stress. This is supported by the finding that treating cells with N-acetyl cysteine during TbTim17 RNAi induction reduced the EC50 of paraquat to wild-type levels. TbTim17 knockdown also increased tolerance of *T. brucei* to heat stress. Either heat or oxidative stress did not increase mitochondrial heat shock protein 70 or Bip levels in the ER in *T. brucei*; instead, they moderately increased TbTim17 levels, which can replenish mitochondrial proteomes. Furthermore, TbTim17 knockdown caused a significant reduction in SL RNA and a 2- to 5-fold increase in tSNAP42 transcript levels, suggesting that mitochondrial stress is linked to the ER stress response pathway in *T. brucei*. Together, these results show that the mitochondrial stress response is primarily mediated by antioxidant defense mechanisms and that TbTim17 plays a protective role in mitochondria under stress in *T. brucei*.

**Importance:** *Trypanosoma brucei*, a parasitic protozoan, is the infectious agent of a deadly disease in humans and livestock known as African trypanosomiasis. TbTim17 is the major component of the TbTIM complex that imports hundreds of nuclear-encoded proteins into the mitochondrial matrix and inner membrane. Here, we show that depletion of TbTim17 induces oxidative stress, which upregulates mitochondrial antioxidant defense mechanisms to mitigate this stress. Subsequently, the SLS response is induced to eliminate the defective parasite from the population. In the wild-type parasite, TbTim17 levels are increased in mitochondria under oxidative and heat stresses, likely to replenish damaged mitochondrial proteomes. This is unlike in other eukaryotes, where oxidative stress degrades Tim17 to induce mitochondrial stress response. Understanding the mitochondrial stress response in *T. brucei* and the role of mitochondrial protein translocases in this process is critical for elucidating the mechanisms of adaptation to environmental stresses and drug resistance in this parasite.

## Introduction

*Trypanosoma brucei*, a divergent eukaryotic parasite, is the causative agent of a fatal disease known as African sleeping sickness in humans and Nagana in ruminants, posing a major health concern and hindering economic development in the affected areas (1). *T. brucei* possesses a single reticular mitochondrion per cell that not only plays a vital role in parasite survival but also modulates its structure and metabolic pattern for adapting under different environmental conditions during parasite differentiation (2). However, mitochondrial stress response pathways are not clearly defined in this organism. Furthermore, trypanosomatids mostly lack transcriptional control of gene expression (3); thus, retrograde signaling from mitochondria to the nucleus to activate the canonical transcription factors is unfound.

As in other eukaryotes, most mitochondrial proteins in *T. brucei* are nucleus-encoded and imported into the mitochondrion from the cytosol (4, 5). *T. brucei* translocates proteins and tRNAs into the mitochondrion via the atypical ATOM and the TbTIM17 complexes of the organellar outer and inner membranes, respectively (6, 7). The TbTIM17 is a mega-Dalton-sized protein complex, consisting of a few relatively conserved proteins and several trypanosome-specific components (6, 8). TbTim17 is the major component of the complex and belongs to the Tim17/22/23 family of proteins (9). In other eukaryotes, Tim17 and Tim23 form the TIM23 complex, whereas Tim22 forms the TIM22 complex (10, 11). In contrast to the 3 proteins in this family in other eukaryotes, *T. brucei* possesses only TbTim17. The Tim17/22/23 family proteins (17-23 kDa) have 4 transmembrane domains centrally located, leaving both the N- and C-terminal hydrophilic regions in the intermembrane space (4, 10, 11). Yeast has one copy each, but multiple copies of these proteins are found in plants and mammals. Humans have two copies of Tim17 (Tim17A and Tim17B) (12, 13). Tim17B is constitutively expressed, whereas Tim17A expression varies across tissues and is upregulated in many forms of cancer (14). It has been shown that Tim17A in humans and Tim23 in *Caenorhabditis elegans* play crucial roles in the mitochondrial unfolded protein response (UPRmt) (15, 16). Mitochondrial stress triggers degradation of Tim17A by the mitochondrial protease YME1L, perturbing mitochondrial import of the transcription factor ATFS-1, which under normal conditions is imported into the mitochondria and quickly degraded by the matrix protease LONP. ATFS-1 possesses a mitochondrial targeting signal (MTS) and a nuclear localization signal (NLS). Thus, unimported ATFS-1 accumulates in the nucleus and activates expression of various chaperones and mitochondrial biogenesis proteins to alleviate the stress (15, 16). The mitochondrial stress response is also connected to the integrated stress response pathways (ISR), which involve many endoplasmic reticulum (ER) and cytosolic proteins that lead to translational attenuation via phosphorylation of the translation initiation factor eIF2-α (17). Besides, there are several other mitochondrial quality control (MQC) pathways. These include mitochondrial precursors overaccumulation stress (mPOS) (19), the constitutively active mitochondrial protein translocation-associated degradation (mitoTAD) (20), and others. In these cases, translocation-impaired proteins are ubiquitinated in the cytosol and degraded by the proteosome.

Stress response pathways have also been studied in trypanosomes. A mostly investigated one is the ER-stress response by splice-leader silencing (SLS-response) (21). Inhibition of protein translocation in the ER or accumulation of the unfolded proteins in the ER-lumen triggers down-regulation of the synthesis of SL-RNA, which is a piece of RNA that needs to be trans-spliced to the 5’-end of each mRNA to make it functional in trypanosomes (3). Thus, SL-RNA silencing reduced the ER protein load and induced apoptotic-like cell death upon consistent stress. Although well-established in the procyclic form that dwells in the insect gut, the existence of the SLS response and an associated upregulation of the ER-chaperone protein, Bip, is controversial in the mammalian infective bloodstream forms of *T. brucei* (22). It is also not clear if a global translation inhibition by eIF2-phosphorylation exists in trypanosomes. Therefore, the ISR pathway is still very sketchy in this parasite. Another study on the MQC pathway showed that a nuclear-localized ubiquitin-like protein (TbUbl1) is released into the cytosol and is required for the degradation of accumulated precursor proteins when mitochondrial protein import is impaired (23). It has also been shown that nutritional stress causes an accumulation of tRNAThr halves in mitochondria, which associate with mitochondrial ribosomes and stimulate translation, thereby increasing *T. brucei* survival (23, 24).

Modulation of mitochondrial activities is critical for *T. brucei* differentiation during its digenetic life cycle. In the mammalian stage, the bloodstream form of *T. brucei* survives by aerobic glycolysis, utilizing glucose as its primary energy source (25). This form lacks cytochrome-dependent respiratory complexes but possesses trypanosome alternative oxidase (TAO) (26). Whereas, upon differentiation to the procyclic form in the insect gut, where glucose is scarce, *T. brucei* utilizes amino acids as its major energy source. At this stage, the mitochondrion is fully developed, with cytochrome-dependent OXPHOS complexes, and TAO levels are reduced. Thus, this form has two terminal oxidases in mitochondria, and the flow of electrons can be flexible. Electron transfer via TAO is non-proton-motive and is not coupled to ATP synthase. It acts as an electron overflow mechanism, thereby reducing mitochondrial oxidative stress (27, 28).

Here, we show that *T. brucei* induces antioxidant defense mechanisms to attenuate mitochondrial stress. Forced downregulation of TbTim17 stimulates this defense mechanism. Furthermore, TbTim17 levels were moderately upregulated under stress, suggesting it has a protective role in mitochondria. Persistent oxidative stress induced by TbTim17 depletion activates the SLS response, leading to cell death.

## Results

### TbTim17 depletion increased *T. brucei* tolerance to paraquat

TbTim17 is the major component of the single TIM complex in *T. brucei* and is essential for the import of hundreds of nuclear-encoded proteins destined to the mitochondrial matrix and inner membrane. To investigate whether TbTim17 depletion elicits a stress response, we induced TbTim17 RNAi in procyclic cells with doxycycline for 48 h. After that, we treated cells with different concentrations of paraquat, which induces mitochondrial oxidative stress, and monitored cell growth for the next 48 h. We used parental (29–13) and TbTim62 RNAi procyclic cells in parallel to compare them. TbTim62 is also a component of the TbTIM17 complex, but its abundance is lower, and the effect of its depletion on the mitochondrial protein import is less than that of TbTim17 (Singha et al., 2012). Results showed that TbTim17-depleted cells are more resistant to paraquat in comparison to the 29-13 and TbTim62-depleted procyclic cells (Fig. 1A). Estimation of the EC50 from three independent experiments showed that the EC50 of paraquat for 29-13 was 100 ± 7.6 μM, whereas that for TbTim17 knockdown cells was more than 300 ± 17.6 μM. Thus, TbTim17 knockdown cells has 3-fold higher EC50 for paraquat in comparison to the 29-13 (Fig. 1B). TbTim62 knockdown cells behave similarly to the 29-13 upon paraquat treatment. These results indicated that TbTim17 depletion equipped *T. brucei* cells to resist paraquat-induced oxidative stress. In other eukaryotes, reduced mitochondrial protein import elicits a stress response to maintain cell homeostasis. Therefore, our results suggest TbTim17 depletion may induce a similar stress response phenomenon in *T. brucei*. Immunoblot analysis of the mitochondrial proteins isolated from cells after 48 h of RNAi induction showed depletion of TbTim17 and TbTim62 levels in respective cell lines, as expected (Fig 1C).

**Figure 1.**
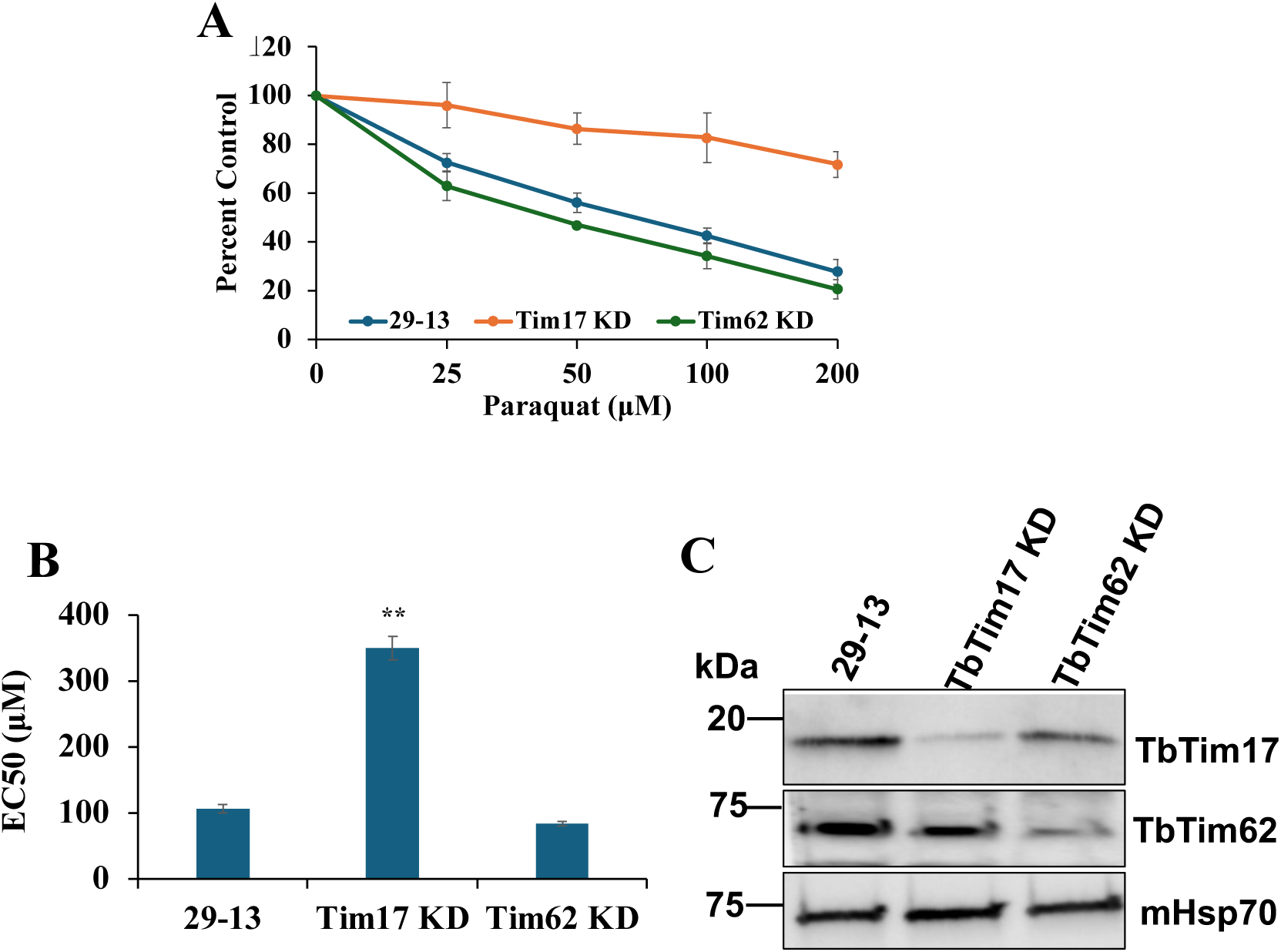
TbTim17 knockdown in *T. brucei* increased EC50 of paraquat. The TbTim17 RNAi and TbTim62 RNAi cell lines were induced with doxycycline for 2 days to knockdown (KD) the target proteins. These cells and the parental control *T. brucei* 29-13 were then treated with different concentrations of paraquat (0-200 μM), and cell growth was monitored for 48 h. **A.** Percent reduction in cell number relative to no paraquat was plotted against different concentrations of paraquat. Standard deviations (SDs) were calculated from 4 independent experiments. **B.** EC50 values of paraquat for 29-13, TbTim17 KD, and TbTim62 KD cells were calculated as described in the materials and methods and plotted. SDs were calculated from 4 independent experiments. Significance values were calculated by a t rest and are indicated by asterisks (**, p <0.05). **C.** Immunoblot analysis of equal amounts of mitochondrial proteins isolated after 48 h of induction of RNAi was performed using antibodies for TbTim17 and TbTim62. *T. brucei* 29-13 cells were grown for the same time and used in parallel. The mHsp70 was used as the loading control.

### Increased ROS due to TbTim17 depletion promotes stress tolerance

It has been shown in mammals and certain nematodes that mitochondrial stress response occurs via retrograde signaling by ATFs transcription factors (15, 16). However, there are few studies on mitochondrial stress response in *T. brucei.* In particular, *T. brucei* lacks transcriptional control of gene expression and any ATFs homolog. To investigate why TbTim17-depleted *T. brucei* is more tolerant to paraquat, we assessed stress markers, such as altered mitochondrial membrane potential and ROS, at the early stages (within 48 h) of TbTim17 RNAi induction. We didn’t observe any significant changes in the mitochondrial membrane potential during this period of TbTim17 RNAi (Fig. 2A). Next, we assessed the levels of ROS in TbTim17-depleted and control cells using DCFH-DA (Fig. 2B). We noticed that the levels of ROS were significantly higher in TbTim17-depleted cells in comparison to 29-13 and TbTim62-depleted *T. brucei*. The 29-13 cells treated with paraquat were used as the positive control. We observed that paraquat treatment increased ROS levels in 29-13 cells, as expected. Therefore, we conclude that TbTim17 knockdown specifically increased mitochondrial ROS levels.

**Figure 2.**
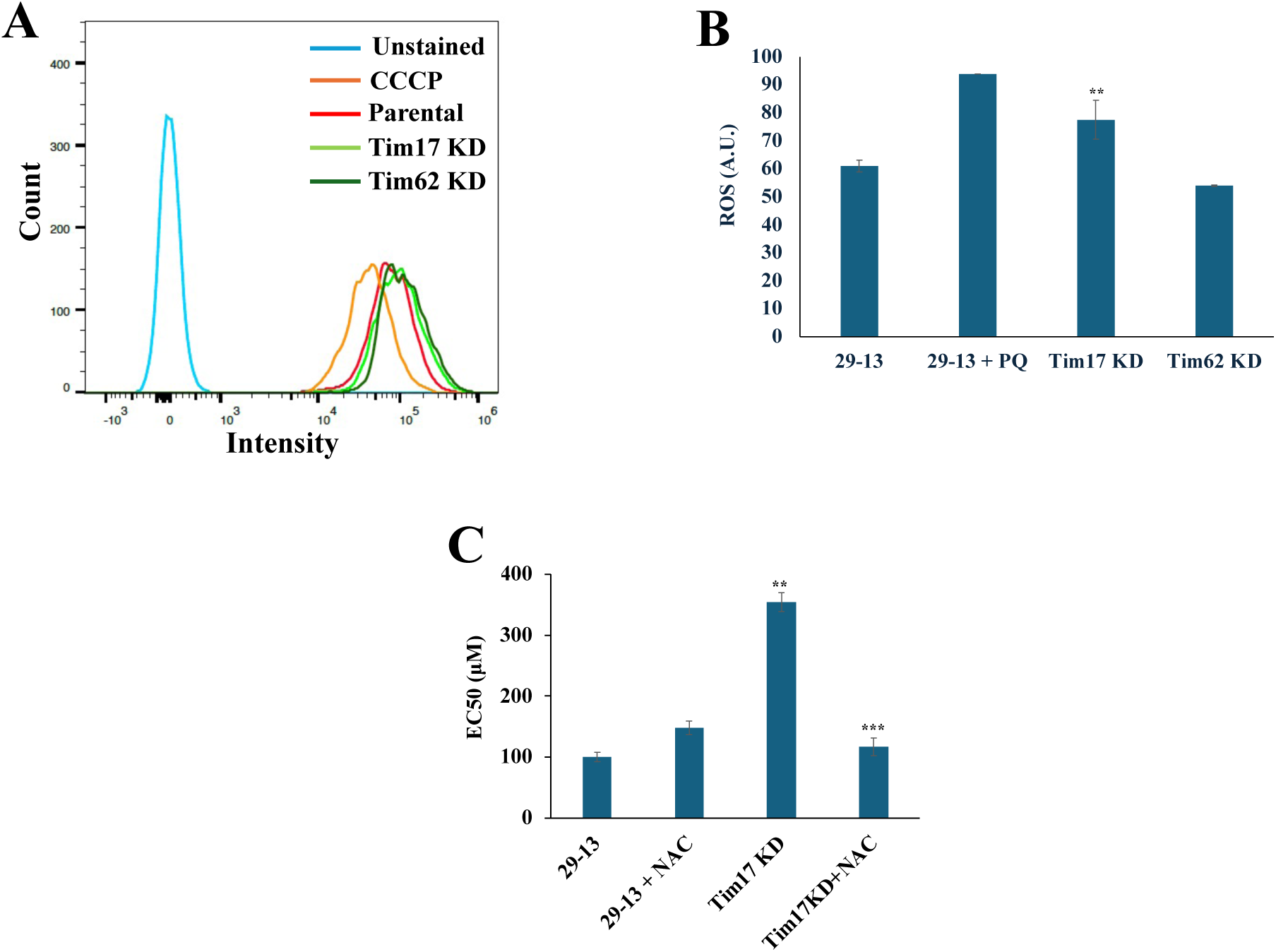
Measurement of mitochondrial membrane potential and ROS levels and treatment of cells with NAC. **A.** *T. brucei* 29-13, TbTim17 RNAi, and TbTim62 RNAi cells induced with doxycycline for 48 h were stained with MitoTracker Red, fixed with 0.37% paraformaldehyde, and fluorescence intensity was measured at 599 nm by fluorescence-activated cell sorter (FACS) analysis. The representative data from multiple experiments are shown. Wild-type cells treated with 50 μM CCCP served as a control to demonstrate mitochondrial membrane potential depletion. **B.** *T. brucei* 29-13, TbTim17 RNAi, and TbTim62 RNAi cells induced with doxycycline for 48 h were stained with DCFH-DA and fluorescence intensity was measured by FACS analysis. Fluorescence intensities were presented in arbitrary units (A.U.). Experiments were done in triplicate, and SDs are shown. **C.** The TbTim17 RNAi cells were induced with doxycycline for 48 h in the absence and presence of NAC (100 μM). *T. brucei* 29-13 cells were also treated with NAC for the same time. EC50 for paraquat was determined as described above and plotted. SDs were calculated from 3 independent experiments. Significance values were calculated (B and C) by a t rest and are indicated by asterisks (**, p <0.05, and ***, p <0.01).

Increased cellular ROS is detrimental to cellular function; however, a certain level of ROS often generates a stress signal that mitigates its effects (29). Therefore, to investigate whether the increased ROS levels in TbTim17 knockdown cells are the cause of these cells becoming more tolerant to paraquat, we treated cells with N-acetyl cysteine (NAC), an antioxidant, during RNAi induction and determined the EC50. The 29-13 cells pretreated with NAC didn’t show any significant changes in the EC50 of paraquat. However, TbTim17 knockdown cells pretreated with NAC showed a significant reduction in EC50 compared with those without NAC pretreatment (Fig. 2C). These results clearly show that increased ROS in TbTim17-depleted cells induces a mitochondrial stress response, possibly by altering the levels of certain stress-tolerant proteins.

### TbTim17 depletion increases SODA levels

We wanted to investigate if TbTim17 depletion induces the expression of chaperones like mitochondrial Hsp70 and Bip, the Hsp70 homolog in the ER, as well as oxidative stress response proteins such as mitochondrial SODA and TAO during 0 to 48 h of induction of TbTim17 RNAi. Results from the immunoblot analysis show that TbTim17 levels decreased gradually with increasing time of RNAi induction, as expected. We didn’t observe any significant increase in the level of mitochondrial Hsp70 (Fig. 3A and B). In fact, mHsp70 levels were reduced slightly at the initial time points but recovered to normal levels by 48 h. In contrast, the SODA levels gradually increased during this period. The TAO level was also elevated at times. However, we observed that the TAO level is also upregulated in control cells during the latter phase of cell growth (Fig. 3A). It was expected that the levels of mitochondrial proteins such as SODA, TAO, and Hsp70 would be reduced due to inhibition of their mitochondrial import in TbTim17 knockdown cells. However, the results indicate that these protein levels remain similar or increase, possibly due to increased protein stability under oxidative stress. The level of Bip, the ER chaperone, remained largely unchanged during this period. These results showed that TbTim17 depletion increased mitochondrial SODA and TAO levels, which is meaningful, as we found that TbTim17 knockdown increased ROS. Together, this suggests that increased ROS due to TbTim17 knockdown acted as a signaling factor, upregulating SOD A and TAO levels, and that these cells became more resistant to paraquat-induced oxidative stress.

**Figure 3.**
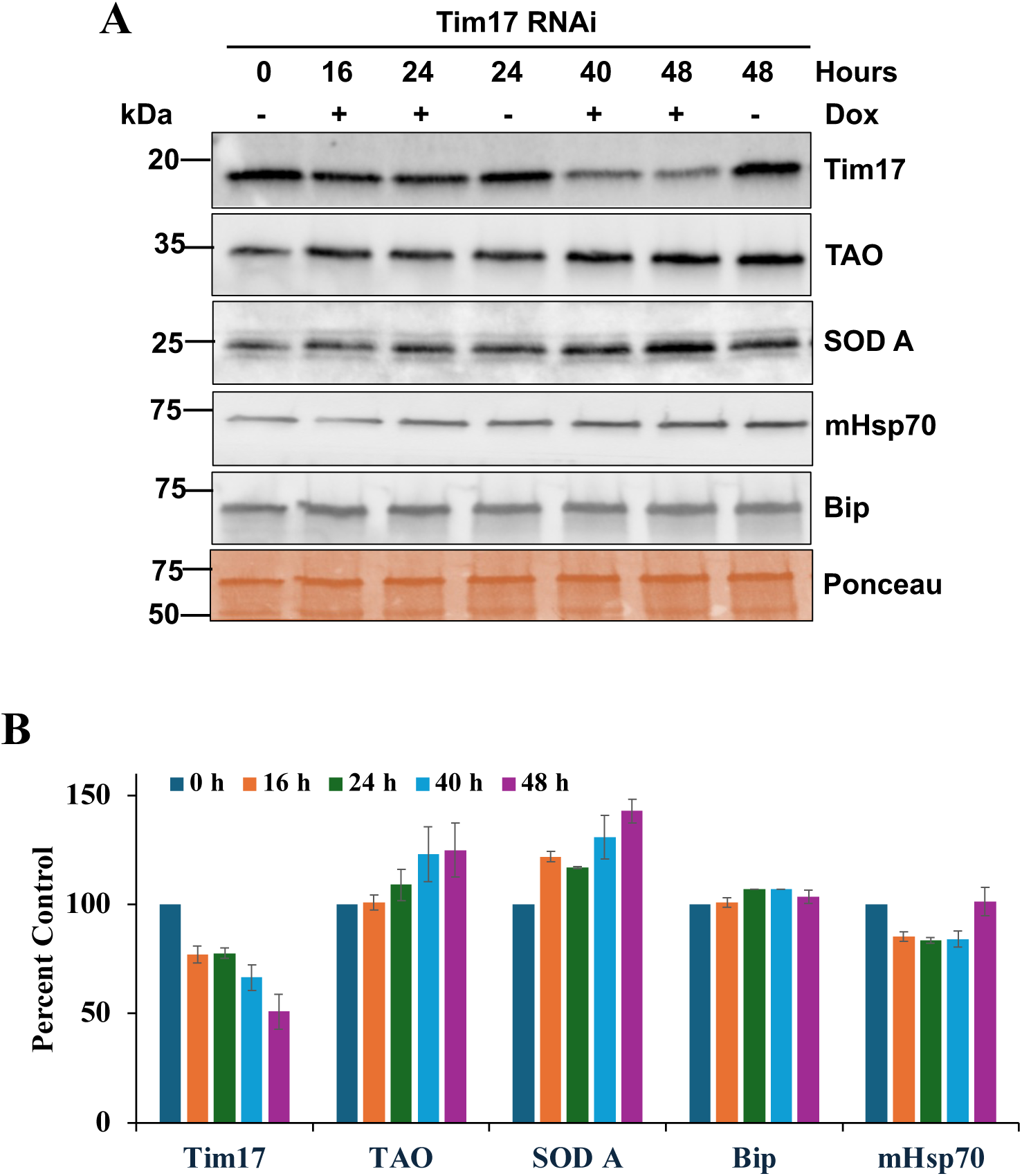
Effect of TbTim17 knockdown on the chaperones and antioxidant proteins in *T. brucei*. **A.** Immunoblot analysis of proteins in the crude membrane fractions isolated from TbTim17 RNAi cells grown in the presence (+) and absence of doxycycline for different times (0-64 h). Blots were probed with the indicated antibodies. Ponceau-stained blot (50 kDa region) is shown as a loading control. **B.** Quantitation of the immunoblot results. Band intensities were quantified in ImageJ, normalized to the corresponding ponceau-stained bands, and plotted as percent control. SDs were calculated from 3 independent experiments.

### TbTim17 knockdown cells are more tolerant to heat stress

*T. brucei* encounters a quick shift in temperature from 27 ℃ to 37 ℃ during its life cycle (1, 2). Thus, to investigate whether TbTim17 plays a role in heat stress, we tested the effect of temperature on the growth of control and TbTim17 knockdown *T. brucei*. After induction of TbTim17 RNAi for 2 days, we split the culture and incubated it separately for 16 h at 27 ℃ and 37 ℃. The 29-13 and TbTim62 RNAi cells were used as controls in parallel. Interestingly, we found that TbTim17 knockdown cells are more tolerant to heat stress than the 29-13 and TbTim62 RNAi cells (Fig. 4A and B). We observed a 10-20 % growth retardation of 29-13 cells due to incubation at 37 ℃ for 16 h (Fig. 4A). In contrast, TbTim17 knockdown cells grew better at 37 ℃ than at 27 ℃. Following TbTim17 knockdown, *T. brucei* cell growth was reduced to 50% compared to the control at 27 ℃. However, after shifting to 37 ℃, these cells grew similarly to control cells at 37 ℃ (Fig. 4A and B). Instead, Tim62 knockdown reduced 15-20% cell growth in comparison to control at 27 ℃, and it is reduced more when transferred at 37 ℃ (Fig. 4A and B). These results clearly show that TbTim17 knockdown cells are better equipped to withstand the stress induced by a temperature increase from 27 ℃ to 37 ℃. Heat stress denatures proteins, forms protein aggregates, and increases ROS levels, accelerating protein denaturation and aggregation and inhibiting cell growth. TbTim17 knockdown cells have higher levels of antioxidant proteins, such as SODA. Therefore, these cells, when transferred at higher temperatures, can mitigate ROS levels that favor cell growth.

**Fig. 4.**
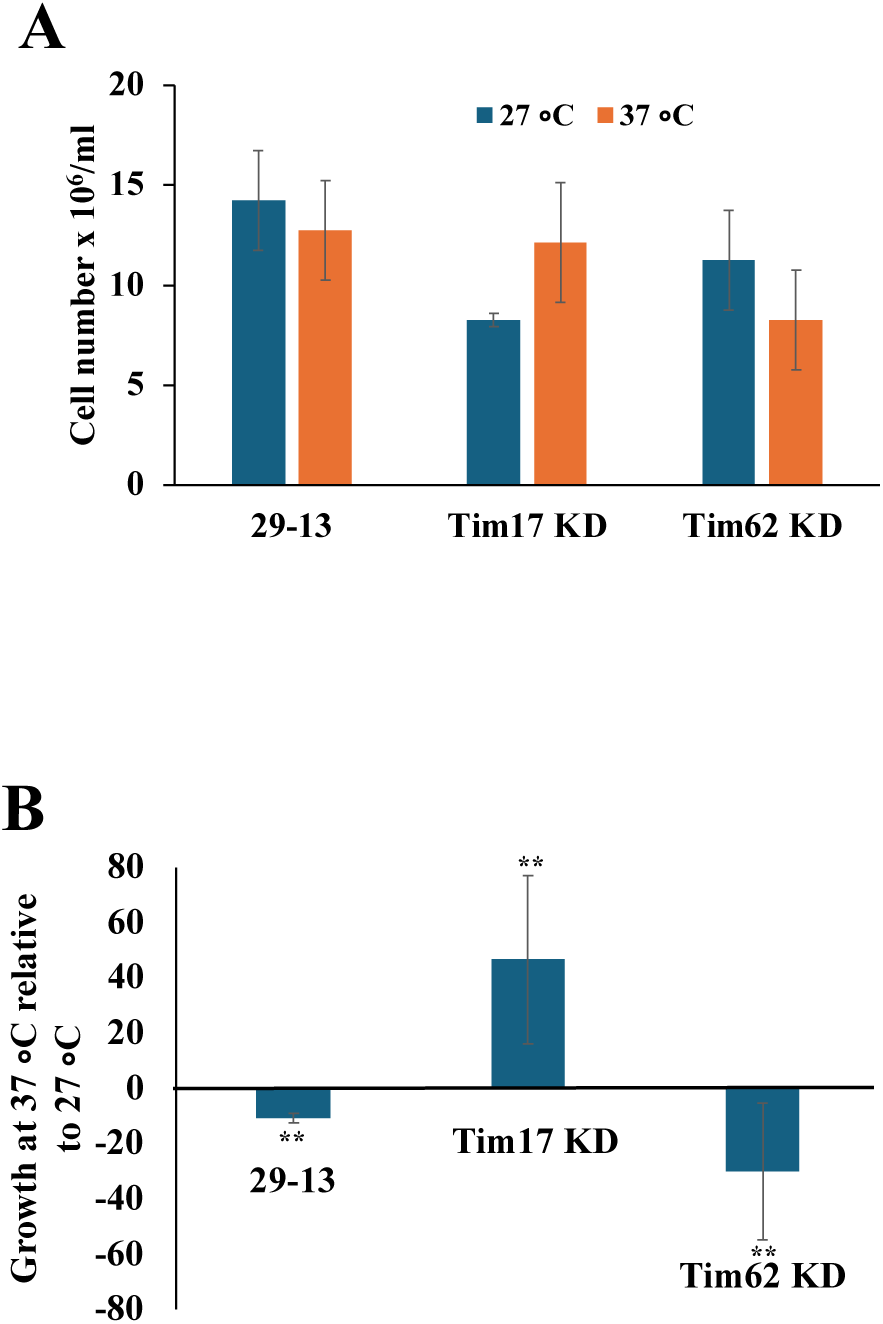
Heat tolerance of the parental and TbTim17 and TbTim62 knockdown *T. brucei*. TbTim17 RNAi and TbTim62 RNAi cells were induced with doxycycline for 48 h and further incubated at different temperatures (27 ℃ and 37 ℃) for 16 h. The parental 29-13 cells were treated similarly for the same period. **A.** Cell numbers were calculated and plotted for each type of cell as indicated. SDs were calculated from three independent experiments. **B.** Cell growth at 37 ℃ relative to that at 27 ℃ for the respective cell types was calculated in percent and plotted. Significance values were calculated by a t rest and are indicated by asterisks (**, p <0.05).

### TbTim17 level is moderately upregulated under stress

Previous reports in other eukaryotes demonstrated that oxidative stress reduced Tim17 levels by inducing its degradation via the mitochondrial protease YME1L (15). Therefore, we sought to determine whether TbTim17 levels are altered under oxidative and heat stress in *T. brucei*. To induce oxidative stress, we treated 29-13 cells with paraquat (100 μM) for shorter (0 to 6 h) and longer (16 to 40 h) time periods and assessed the levels of TbTim17, TAO, SOD A, mHsp70, and Bip. We found that, upon treatment with paraquat, TbTim17 levels are upregulated over time, as are TAO and SODA (Fig. 5A and B). Paraquat generates ROS by perturbing the cytochrome-dependent respiratory pathway (30). Thus, the TAO level was upregulated upon paraquat treatment to increase respiration through the alternative pathway. In contrast, mHsp70 and Bip were not upregulated. Together, these results suggest that TbTim17 level is increased to protect mitochondria under stress. Similarly, to investigate if TbTim17 levels were upregulated at higher temperature, we compared TbTim17 levels in 29-13, TbTim17 RNAi, and TbTim62 RNAi cells incubated at 27 ℃ and 37 ℃. We observed that TbTim17 levels were consistently higher in 29-13 cells grown at 37 ℃ than in those grown at 27 ℃ (Fig. 5C and D). We observed that TAO levels were slightly reduced due to heat stress across all three cell types. Since TAO is thermogenic, reduced respiration via TAO could be a compensatory mechanism at higher temperatures. SODA levels were slightly reduced upon heat treatment in 29-13 cells as reported previously (); however, they stayed up at both temperatures in RNAi cells. In consistency with the effect of oxidative stress, there were no significant changes in mHsp70 levels under heat stress.

**Fig. 5.**
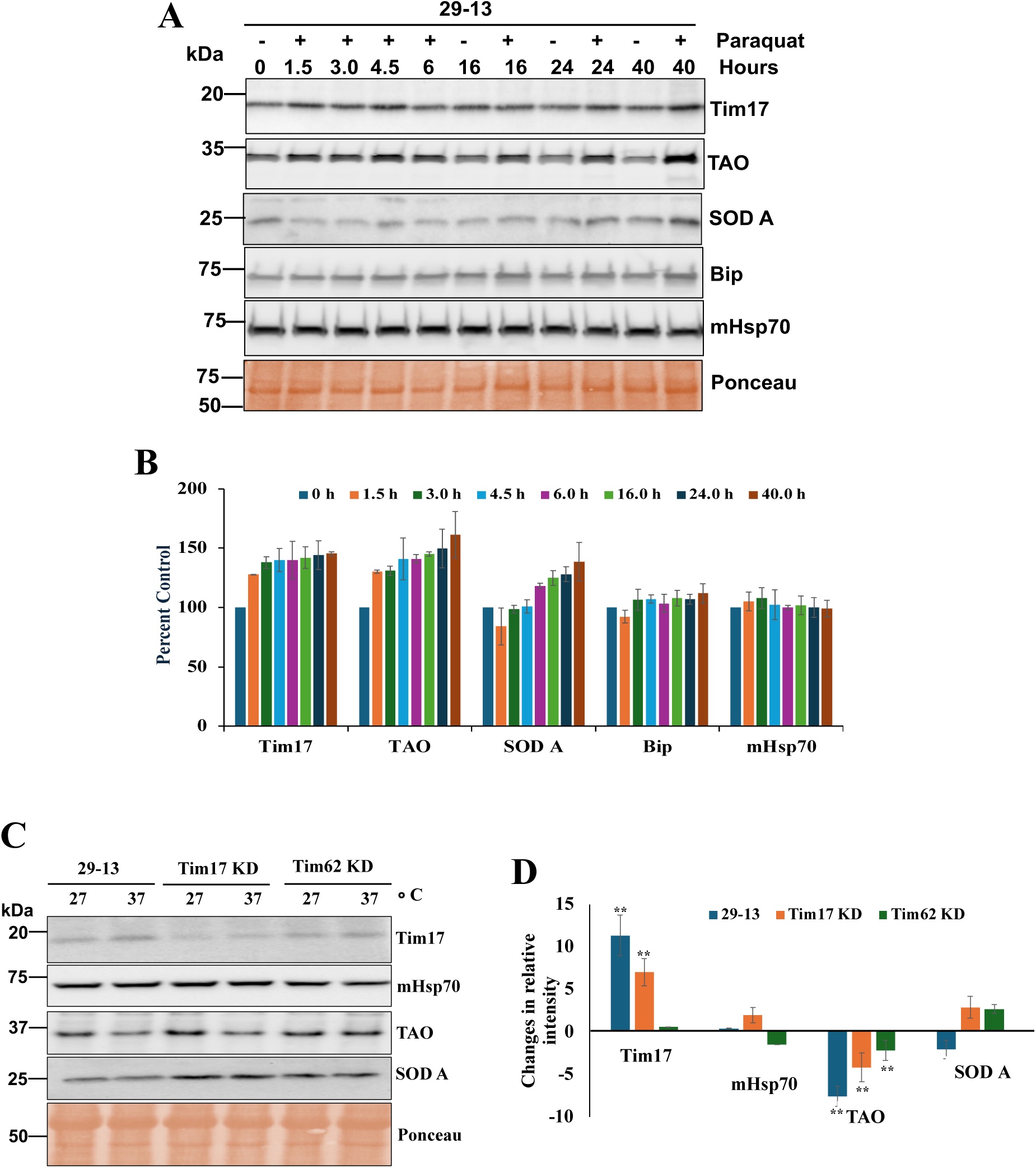
Effect of paraquat treatment on the chaperones and antioxidant proteins in *T. brucei*. **A.** Immunoblot analysis of proteins in the crude membrane fractions isolated from *T. brucei* 29-13 cells treated (+) and kept untreated (−) for different time points (0-40 h). Blots were probed with the indicated antibodies. Ponceau-stained blot (50 kDa region) is shown as a loading control. **B.** Quantitation of the immunoblot results. Band intensities were quantified in ImageJ, normalized to the corresponding ponceau-stained bands, and plotted as percent control. SDs were calculated from 3 independent experiments. **C.** TbTim17 RNAi and TbTim62 RNAi cells were induced with doxycycline for 48 h and further incubated at different temperatures (27 ℃ and 37 ℃) for 16 h. The parental 29-13 cells were treated similarly for the same period. Cells were harvested, and total cellular proteins were analyzed by immunoblot using the indicated antibodies. **D.** Band intensities were quantified by ImageJ software, normalized with the corresponding ponceau-stained bands. Changes in the intensity of each protein band at 37 ℃ relative to that at 27 ℃ in each cell type were calculated as percentages and plotted. SDs were calculated from 3 independent experiments.

### TbTim17 knockdown induces SLS response

One specific stress response phenomenon in *T. brucei* is the SLS response, which is activated upon ER stress. During this response, the SL RNA transcription is blocked by upregulation of the tSNAP42, a specific transcription factor for SL RNA synthesis. To examine if TbTim17 RNAi could induce the SLS response, we quantified the transcript levels of SL RNA and tSNAP42 in 29-13 and TbTim17 knockdown *T. brucei* by qRT-PCR. We found ∼50% reduction of SL-RNA and 2 to 5-fold increase in the level of tSNAP42 transcript in TbTim17 knockdown cells in comparison to the control (Fig. 6A). These results indicate that inhibition of mitochondrial protein import by TbTim17 knockdown induces SLS response to reduce the protein load in the mitochondria.

**Fig. 6.**
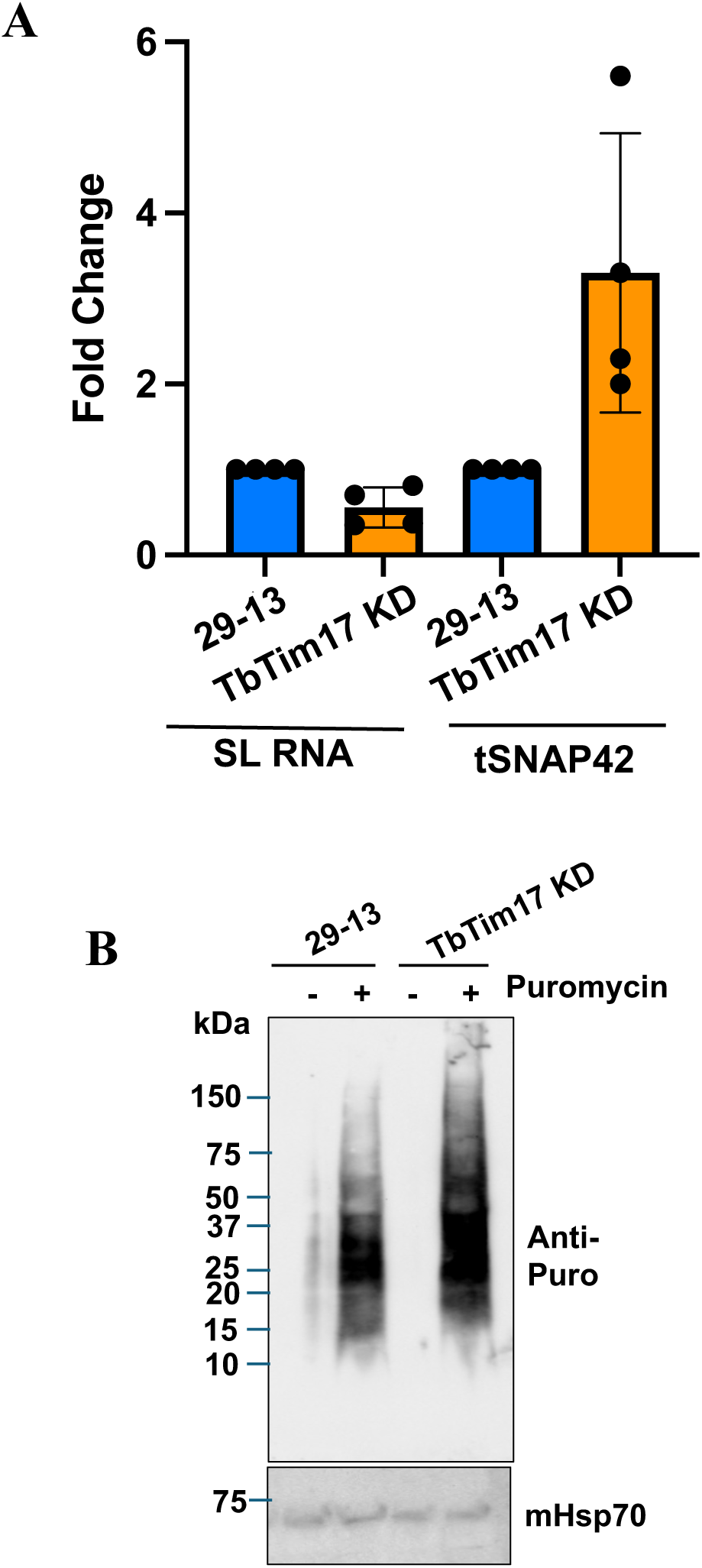
Effect of TbTim17 RNAi on the levels of SL-RNA and tSNAP42 transcripts. **A.** qRT-PCR analyses were performed to estimate the levels of SL-RNA and tSNAP42 transcripts. Total RNA was isolated from *T. brucei* 29-13 and TbTim17 RNAi cells induced with doxycycline for 48 h. cDNA was synthesized from DNase-treated RNA, and qRT-PCR was performed as described in the Materials and Methods. SL RNA levels were normalized to 7SL, and tSNAP42 transcript levels were normalized to actin. Results from four independent biological replicates are presented. **B.** Assay for puromycin incorporation in newly synthesized proteins. *T. brucei* 29-13 and TbTim17 RNAi cells induced with doxycycline for 48 h and were incubated with puromycin (2 μg/ml) for 1 h. Cells were washed, and total cellular proteins were analyzed by SDS-PAGE and immunoblot analysis probed with anti-puromycin antibody.

To examine if TbTim17 depletion induces a global translation inhibition, we performed a puromycin incorporation assay. Puromycin acts as an analog of tyrosyl tRNA, enters the ribosomes’ A-site, covalently binds to the growing peptide chain, and causes premature termination. Therefore, puromycin incorporation into protein can be used to measure active translation in cells (30). Incubation of the 29-13 and TbTim17 knockdown cells in the presence of puromycin and analysis of the total cellular proteins by immunoblot using anti-puromycin antibody, we found that incorporation of puromycin in the newly synthesized proteins are very similar to both types of cells, showing that stresses induced by TbTim17 RNAi for 48 h didn’t induce a global translation inhibition (Fig. 6B). These results were not unexpected as it has been shown that *T. brucei* does not respond to stress by translational inhibition mediated by eIF2 phosphorylation like in other eukaryotes (31). Instead, SLS response induces apoptotic-like cell death in *T. brucei* when the stress persists (21). Therefore, TbTim17 knockdown increases antioxidant defense mechanisms to mitigate stress and ultimately induces an SLS response that proceeds to apoptotic-like cell death.

## Discussion

Here, we demonstrated that TbTim17, a major and essential component of the mitochondrial protein import machinery, plays a role in the mitochondrial stress response; however, it acts differently than its counterparts in other eukaryotes. Depletion of TbTim17 increased ROS levels within 24-48 h, which could be due to a reduction in the import of multiple nuclear-encoded mitochondrial proteins simultaneously or to the inhibition of the import of specific labile protein(s) needed to maintain ROS levels. Consequently, mitochondrial Fe-SODA level was upregulated. SODA converts ROS to hydrogen peroxide. Trypanosomes lack catalase but possess a strong peroxidase system, including tryparedoxin peroxidase (TXN) and glutaredoxin 1 (GrX1) in mitochondria (32), which neutralize peroxides via a glutathione/trypanothione redox transfer chain to maintain cell homeostasis. SODA is a nuclear-encoded protein that is required to be imported to the mitochondrial matrix via the translocase system. Therefore, upregulation of SODA protein levels in TbTim17-depleted cells suggests that this protein may be stabilized under these conditions, possibly via post-translational modification. Furthermore, we showed that the increased ROS levels are responsible for the oxidative stress tolerance of TbTim17-depleted cells. This phenomenon fits well with the increased SODA levels; however, how mitochondria sense the ROS levels to upregulate the levels of SODA requires further investigation. Previous studies showed that the SODA transcript level is upregulated in the G1 and G2 phases of the cell cycle (33), and SODA plays a role in *T. brucei* life-cycle adaptation process and virulence (34, 35). Therefore, there must be specific mechanisms for its regulation in the parasite.

TAO expression is also very flexible. TAO is significantly more abundant in the bloodstream forms than in the procyclic form (26). It is upregulated in the metacyclic form and is modulated under certain experimental and environmental changes (2), suggesting involvement in adaptive or regulatory functions pertinent to survival and life-cycle progression. TAO by-passes electron flow and reduces ROS levels during perturbations of electron flow through the cytochrome-dependent respiratory pathway. We observed a clear upregulation of TAO in the 29-13 procyclic cells treated with paraquat. TAO level was also moderately upregulated in TbTim17 knockdown cells, likely to reduce oxidative stress.

TbTim17 is the single homolog of Tim17/22/23 family proteins in trypanosomes. Native gel electrophoresis revealed that TbTim17 forms multiple-sized protein complexes by associating with several other trypanosome-specific Tim proteins, like TbTim62 (6, 36). A previous report showed that TbTim62 knockdown reduced the abundance of the TbTIM17 complex and the steady-state levels of TbTim17 (37). However, TbTim62 depletion did not increase ROS levels, unlike TbTim17 RNAi within 2 days. This indicates that TbTim17 RNAi reduces the levels of the TbTim17, the direct target of the RNAi, below a certain threshold at this time point, that is not reached by the secondary effect of TbTim62 RNAi.

In addition to oxidative stress, TbTim17-depleted *T. brucei* was more adaptable to a change in temperature from 27 ℃ to 37 ℃. It is speculated that this tolerance is also due to elevated SODA levels. However, we couldn’t eliminate the existence of other mechanisms. Interestingly, we found that TbTim17 levels were slightly upregulated in the 29-13 procyclic cells under both heat and oxidative stresses. This may be a response to upregulate mitochondrial proteomes under stress, when mitochondrial proteins are damaged by oxidation. Tim17 is known to protect mitochondrial DNA in other eukaryotes (38), although the mechanism underlying this function remains unknown. Whether TbTim17 has a similar function is worth investigating. During heat stress, we observed downregulation of TAO in all *T. brucei* cell lines tested, which is also very interesting and likely correlated with TAO’s thermogenic property.

Organellar communication is critical to maintain cellular homeostasis. In this regard, communication between the ER and mitochondria is well established. In *T. brucei*, perturbation of ER protein translocation induces an SLS response by activating the ER-associated kinase, PK3, which, in turn, phosphorylates the TATA-binding protein, TRF4, in the nucleus, thereby silencing SL-RNA transcription (39). It has recently been shown that depletion of TbRHOM1, a TbTim17-associated protein, induces an SLS response to a lesser extent than does depletion of Sec63, the ER translocase (40). This shows that stresses induced by mitochondrial protein import inhibition are transmitted to the ER. This report also showed that depletion of golgi-localized quiescin sulfhydryl oxidase (QSOX) and ER oxidoreductin by RNAi induces the SLS response (40). Here, we further showed that the depletion of TbTim17 reduces the levels of SL-RNA transcripts by about 50% compared with the control. TbTim17 reduction also increased the levels of tSNAP42 transcript by about 4- to 5-fold. This indicates that stresses induced by TbTim17 depletion in mitochondria are transmitted to the ER and induce SLS response. Apparently, this SL RNA silencing had no significant effect on overall protein translation. This is consistent with the previous observation that the SLS response induces an apoptosis-like phenomenon in the procyclic form of *T. brucei* to eliminate unfit cells from the population (21).

Altogether, our results showed that TbTim17 plays a protective role in mitochondria. Heat or oxidative stress consistently increased TbTim17 levels likely to replenish mitochondrial proteomes. TbTim17 depletion induces an antioxidant defense mechanism to mitigate stress caused by mitochondrial protein import inhibition, as we found that these cells were more tolerant to both heat and oxidative stresses. However, TbTim17 depletion is detrimental because it induces cell apoptosis by activating the SLS-response. Therefore, TbTim17 is a potential therapeutic target for African trypanosomiasis.

## Materials and Methods

### Cell lines, growth medium, and cell growth analysis

The procyclic form of *T. brucei 427* 29-13 expressing T7 polymerase and tetracycline repressor genes was grown in SDM-79 medium containing 10% heat inactivated fetal bovine serum in the presence of hygromycin (50 μg/ml), G418 (15 μg/ml) (41) and was used as the parental control for our experiments. *T. brucei* cells expressing TbTim17 RNAi (TbTim17 KD) and TbTim62 RNAi (TbTim62 KD) were previously developed (42, 37). These cell lines were maintained in the same medium supplemented with phleomycin (2.5 μg/ml). Cell growth was assessed by inoculating the procyclic form at a cell density of 2 × 10^6^/ml in fresh medium. Cells were counted at different growth time points with a Neubauer hemocytometer. The log of the cumulative cell number was plotted against incubation times.

### Subcellular fractionation and crude mitochondria isolation

*T. brucei* mid-log phase cells (2 × 10^8^) were resuspended in 500 μl of SMEP buffer (250 mM sucrose, 20 mM morpholinepropanesulfonic acid [MOPS]-KOH, pH 7.4, 2 mM EDTA, 1 mM phenylmethylsulfonyl fluoride [PMSF]) containing 0.03% digitonin and incubated on ice for 5 min. The cell suspension was then centrifuged for 5 min at 6,800 × *g* at 4 °C. The resultant pellet was considered the crude mitochondrial fraction, and the supernatant contained soluble cytosolic proteins.

### Paraquat treatment and EC50 determination

For determination of the growth inhibitory concentration (EC50) of paraquat, mid-log phase procyclic form cells were inoculated in fresh medium as described above in 24-well plates. Calculated amounts of paraquat (stock of 20 mM in DMSO) were added to the culture to reach the final concentrations 0 to 200 μM. Maximum concentration of DMSO didn’t exceed 0.01%. Triplicate wells were set up for each concentration. Cells were counted each day for 2 days from each well twice. EC50 was determined from normalized non-linear regression plots for the percent inhibition of cell growth versus log of paraquat concentrations using GraphPad Prism (version 10). Standard deviations were calculated from triplicates.

### Measurement of cellular ROS

*T. brucei* cells (1 × 10^7^/ml) were incubated with 10 μM 2’,-7’-dichlorodihydrofluorescein diacetate (DCFH-DA) for 30 min at 27 °C. After incubation, cells were washed once with 1 X PBS and analyzed by flow cytometry, by measuring the fluorescence emission at 520 nm using the excitation wavelength at 492 nm.

### Puromycin incorporation assay

Log-phase cells were incubated in the presence and absence of puromycin (2 μg/ml) in complete medium for 1 hour. Cells were harvested by centrifugation, washed 3 times with 1 X PBS and lysed with SDS-PAGE sample buffer. Total cellular proteins were analyzed by SDS-PAGE and immunoblot analyses using anti-puromycin antibody as the probe.

### SDS-PAGE and Immunoblot analysis

Proteins from isolated mitochondria and total cell extracts were resolved on a 12% SDS/PAGE gel and then transferred to nitrocellulose membranes (Bio-Rad). The antigens were visualized by using a West Pico Chemiluminescent Substrate (Thermo Fisher). Antibodies against TbTim17 (42), TbTim62 (37), SODA (43), TAO (44), HSP70 (45), TbBip (46) were used as probes. The secondary antibodies used were either anti-rabbit or anti-mouse immunoglobulins linked to horseradish peroxidase (Thermo Fisher).

### qRT-PCR

For quantitative real-time PCR (qRT-PCR) analysis, RNA was isolated from *T. brucei* procyclic cells using Trizol reagent (Thermo Fisher) according to the manufacturer’s protocol. RNA was treated with RNase-Free Turbo DNase (Thermo Fisher) at 37 ℃ for 30 min and RNA was recovered by phenol/chloroform extraction and ethanol precipitation. The cDNA was made from 1.0 μg of RNA(DNase-digested) using the iScript cDNA synthesis kit (Bio-Rad). Real-time quantitative PCR was performed with a CFX96 Touch real-time PCR detection system (Bio-Rad). The resulting cDNA was amplified using SsoAdvanced universal SYBR green supermix (Bio-Rad) and gene specific primers targeting the following transcripts: spliced leader RNA (nt 26 to 108), 7SL RNA, gene ID Tb927.8.2861 (nt 147 to 221), TbSNAP42, gene ID Tb927.5.3910 (nt 703 to 811), and actin A, gene ID Tb927.9.8850 (nt 1031 to 1091), as described (22). Spliced leader RNA transcripts were normalized to 7SL RNA, while TbSNAP42 transcript levels were normalized to Actin level. All reactions were performed in technical triplicate, and standard deviations were calculated from three biological replicates.

### Densitometry and statistical analyses

ImageJ software (NIH) was used to perform densitometry of Western blots. The band intensity of the loading control (ponceau stained band around 50 kDa region) was used for normalization, and percent control values were plotted using MS Excel. Standard deviations were calculated from three independent experiments for each blot.

## Acknowledgments

This work was supported by the NIH grant 1RO1AI125662 and NSF-2401909 to MC. The Molecular Biology and the Flow-cytometry Core Facility is supported by NIH grant U54RR026140/U54MD007593. We thank George Cross for *T. brucei* cell lines, Norma Andrews for SODA, Paul Englund for the mHsp70, and Jay Bangs for the Bip antibodies.

## References

1. Sternberg JM, Maclean L. 2010. A spectrum of disease in human African trypanosomiasis: the host and parasite genetics of virulence. Parasitology 137:2007–2015

2. Zikova A. 2022. Mitochondrial adaptations throughout the Trypanosoma brucei life cycle. J. Euk. Microbiol.69:e12911

3. Clayton CE. 2016. Gene expression in kinetoplastids. Curr. Opin. Microbiol. 32:46–51

4. Chaudhuri M, Darden C, Gonzalez FS, Singha UK, Quinones L, Tripathi A. 2020. Tim17 updates: A comprehensive review of an ancient mitochondrial protein translocator. Biomolecules 10:1–20.

5. Schneider A. 2020. Evolution of mitochondrial protein import – lessons from trypanosomes. Biol Chem 401:663–676

6. Singha UK, Hamilton V, Duncan MR, Weems E, Tripathi MK, Chaudhuri M. 2012. The Translocase of Mitochondrial Inner Membrane in *Trypanosoma brucei*. J Biol Chem 287:14480–14493

7. Tschopp F, Charrière F, Schneider A. 2011. In vivo study in *Trypanosoma brucei* links mitochondrial transfer RNA import to mitochondrial protein import. EMBO Rep. 12:825–832

8. Harsman A, Oeljeklaus S, Wenger C, Huot JL, Warscheid B, Schneider A. 2016. The Non-canonical mitochondrial inner membrane presequence translocase of trypanosomatids contains two essential rhomboid-like proteins. Nat. Commun 7:13707.

9. Weems E, Singha UK, Hamilton V, Smith JT, Waegemann K, Mokranjac D, Chaudhuri M. 2015. Functional complementation analyses reveal that the single PRAT-family protein of *Trypanosoma brucei* is a divergent homolog of Tim17 in *Saccharomyces cerevisiae*. Eukaryot Cell. 14:286–296.

10. Herrmann JM, Bykov Y. 2023. Protein translocation in mitochondria: Sorting out Toms, Tims, Pams, Sams and Mia. FEBS Lett 597:1553–1554

11. Busch JD, Fielden LF, Pfanner N, Wiedemann N. 2023. Mitochondrial protein transport: versatality of translocases and mechanisms. Mol. Cell. 83:890–910

12. Bauer MF, Gempel K, Reichert AS, Rappold GA, Lichtner P, Gerbitz KD, Neupert W, Brunner M, Hofmann S.1999. Genetic and structural characterization of the human mitochondrial inner membrane translocase, J. Mol. Biol. 289:69–82

13. Sinha D, Srivastava S, Krishna L, D’Silva P. 2014. Unraveling the intricate organization of mammalian mitochondrial presequence translocases: existence of multiple translocases for maintenance of mitochondrial function. Mol. Cell. Biol. 34:1757–75

14. Chan AA, Sankar K, Reckamp KL, Díaz B, Lee DJ. 2025. Elevated co-expression of TIMM17A and NMT1 is associated with poor survival in non-small cell lung cancer. Sci. Rep. 15:35597

15. Rainbolt TK, Atanassova N, Genereux JC, Wiseman RL. 2013. Stress-regulated translational attenuation adapts mitochondrial protein import through Tim17A degradation. Cell Metab. 18:908–919.

16. Nargund AM, Pellegrino MW, Fiorese CJ, Baker BM, Haynes CM. 2012. Mitochondrial import efficiency of ATFS-1 regulates mitochondrial UPR activation. Science.337:587–590

17. Genuth NR, Dillin A. 2025. Translational regulation in stress biology. Nat. Cell. Biol. 27:1609–1621

18. Wang X, Chen XJ. 2015. A cytosolic network suppressing mitochondria-mediated proteostatic stress and cell death. Nature 524:481–484

19. Mårtensson CU, Priesnitz C, Song J, Ellenrieder L, Doan KN, Boos F, Floerchinger A, Zufall N, Oeljeklaus S, Warscheid B, Becker T. 2019. Mitochondrial protein translocation-associated degradation. Nature. 569:679–683

20. Michaeli S. 2012. Spliced leader RNA silencing (SLS) - a programmed cell death pathway in Trypanosoma brucei that is induced upon ER stress. Parasite. Vector, 5:107.

21. Tiengwe, C., Brown, A. E., & Bangs, J. D. 2015. Unfolded Protein Response Pathways in Bloodstream-Form *Trypanosoma brucei*. Euk. Cell, 14:1094–1101.

22. Fricker R, Brogli R, Luidalepp H, Wyss L, Fasnacht M, Joss O, Zywicki M, Helm M, Schneider A, Cristodero M, Polacek N. 2019. A tRNA half modulates translation as stress response in *Trypanosoma brucei*. Nat. Commun. 10:118

23. Brogli, R., Cristodero, M., Schneider, A., & Polacek, N. 2023. A ribosome-bound tRNA half stimulates mitochondrial translation during stress recovery in *Trypanosoma brucei*. Cell Rep. 42:113112.

24. Verner, Z., Basu, S., Benz, C., Dixit, S., Dobáková, E., Faktorová, D., Hashimi, H., Horáková, E., Huang, Z., Paris, Z., Peña-Diaz, P., Ridlon, L., Týč, J., Wildridge, D., Zíková, A., & Lukeš, J. 2015. Malleable mitochondrion of *Trypanosoma brucei*. Int. Rev.Cell mol. Biol. 315:73–151.

25. Chaudhuri, M., Ott, R. D., & Hill, G. C. 2006. Trypanosome alternative oxidase: from molecule to function. Trends Parasitol. 22:484–491.

26. Fang, J., & Beattie, D. S. 2003. Alternative oxidase present in procyclic Trypanosoma brucei may act to lower the mitochondrial production of superoxide. Arch. Biochem and biophy. 414:294–302.

27. Tsuda A, Witola WH, Konnai S, Ohashi K, Onuma M. 2006. The effect of TAO expression on PCD-like phenomenon development and drug resistance in *Trypanosoma brucei*. Parasitol Int. 55:135–42

28. Aparicio-Trejo, O. E., Hernández-Cruz, E. Y., Reyes-Fermín, L. M., Ceja-Galicia, Z. A., & Pedraza-Chaverri, J. 2025. The role of redox signaling in mitochondria and endoplasmic reticulum regulation in kidney diseases. Arch. Toxico, 99:1865–1891.

29. Wilkinson, S. R., Prathalingam, S. R., Taylor, M. C., Ahmed, A., Horn, D., & Kelly, J. M. 2006. Functional characterisation of the iron superoxide dismutase gene repertoire in *Trypanosoma brucei*. Free Rad Biol Med. 40:198–209.

30. Cabrera-Cabrera, F., Tull, H., Capuana, R., Kasvandik, S., Timmusk, T., & Koppel, I. 2023. Cell type-specific labeling of newly synthesized proteins by puromycin inactivation. J. Biol. Chem. 299:105129.

31. Avila, C. C., Peacock, L., Machado, F. C., Gibson, W., Schenkman, S., Carrington, M., & Castilho, B. A. 2016. Phosphorylation of eIF2α on Threonine 169 is not required for Trypanosoma brucei cell cycle arrest during differentiation. Mol Biochem. Parasitol. 205:16–21.

32. Leroux AE, Krauth-Siegel RL. 2016. Thiol redox biology of trypanosomatids and potential targets for chemotherapyt. Mol. Biochem. Parasitol. 206:67–74

33. Archer SK, Inchaustegui D, Queiroz R, Clayton C. 2011. The cell cycle regulated transcriptome of *Trypanosoma brucei*. PLoS One.6:e18425

34. Mittra, B., Laranjeira-Silva, M. F., Miguel, D. C., Perrone Bezerra de Menezes, J., & Andrews, N. W. 2017. The iron-dependent mitochondrial superoxide dismutase SODA promotes *Leishmania* virulence. J. Biol. Chem. 292:12324–12338.

35. Olmo F, Urbanová K, Rosales MJ, Martín-Escolano R, Sánchez-Moreno M, Marín C. 2015. An in vitro iron superoxide dismutase inhibitor decreases the parasitemia levels of *Trypanosoma cruzi* in BALB/C mouse model during acute phase. Int. J. Parasitol Drugs Drug Resist. 5:110–6

36. Weems, E., Singha, U. K., Smith, J. T., & Chaudhuri, M. 2017. The divergent N-terminal domain of Tim17 is critical for its assembly in the TIM complex in *Trypanosoma brucei*. Mol. Biochem. Parasitol. 218:4–15.

37. Singha, U. K., Hamilton, V., & Chaudhuri, M. 2015. Tim62, a Novel Mitochondrial Protein in *Trypanosoma brucei*, Is Essential for Assembly and Stability of the TbTim17 Protein Complex. J. Biol. Chem. 290: 23226–23239.

38. Matta, S. K., Pareek, G., Bankapalli, K., Oblesha, A., & D’Silva, P. 2017. Role of Tim17 Transmembrane Regions in Regulating the Architecture of Presequence Translocase and Mitochondrial DNA Stability. Mol. Cell. Biol. 37:e00491–16.

39. Hope, R., Egarmina, K., Voloshin, K., Waldman Ben-Asher, H., Carmi, S., Eliaz, D., Drori, Y., & Michaeli, S. 2016. Transcriptome and proteome analyses and the role of atypical calpain protein and autophagy in the spliced leader silencing pathway in *Trypanosoma brucei*. Mol. Microbiol. 102: 1–21.

40. Okalang, U., Mualem Bar-Ner, B., Rajan, K. S., Friedman, N., Aryal, S., Egarmina, K., Hope, R., Khazanov, N., Senderowitz, H., Alon, A., Fass, D., & Michaeli, S. 2021. The Spliced Leader RNA Silencing (SLS) Pathway in Trypanosoma brucei Is Induced by Perturbations of Endoplasmic Reticulum, Golgi Complex, or Mitochondrial Protein Factors: Functional Analysis of SLS-Inducing Kinase PK3. mBio. 12:e0260221.

41. Wirtz E, Hoek M, Cross GA. 1998. Regulated processive transcription of chromatin by T7 RNA polymerase in *Trypanosoma brucei*. Nucleic Acids Res. 26:4626–4634

42. Singha, U. K., Peprah, E., Williams, S., Walker, R., Saha, L., & Chaudhuri, M. 2008. Characterization of the mitochondrial inner membrane protein translocator Tim17 from *Trypanosoma brucei*. Mol. Biochem. Parasitol. 159:30–43.

43. Saldivia, M., Ceballos-Pérez, G., Bart, J. M., & Navarro, M. 2016. The AMPKα1 Pathway Positively Regulates the Developmental Transition from Proliferation to Quiescence in Trypanosoma brucei. Cell Rep. 17:660–670.

44. Chaudhuri M, Ajayi W, Hill GC. 1998. Biochemical and molecular properties of the Trypanosoma brucei alternative oxidase. Mol. Biochem. Parasitol. 95:53–68

45. Effron, P. N., Torri, A. F., Engman, D. M., Donelson, J. E., & Englund, P. T. 1993. A mitochondrial heat shock protein from Crithidia fasciculata. Mol. Biochem. Parasitol. 59:191–200.

46. Bangs, J. D., Uyetake, L., Brickman, M. J., Balber, A. E., & Boothroyd, J. C. 1993. Molecular cloning and cellular localization of a BiP homologue in Trypanosoma brucei. Divergent ER retention signals in a lower eukaryote. J. Cell Sci. 105:1101–1113.

